# Sex-specific calibration of memory recall by glucocorticoid receptors on cortical astrocytes

**DOI:** 10.1101/2020.09.01.278556

**Authors:** WW Taylor, BR Imhoff, ZS Sathi, KM Garza, BG Dias

## Abstract

Dysfunctions in memory recall lead to pathological fear; a hallmark of trauma-related disorders, like Post-Traumatic Stress Disorder (PTSD). Heightened recall of an association between a cue and trauma, as well as impoverished recall that a previously trauma-related cue is no longer a threat both result in a debilitating fear toward the cue. Glucocorticoid-mediated action via the glucocorticoid receptor (GR) influences memory recall. This literature has primarily focused on GRs expressed in neurons or ignored cell-type specific contributions. To ask how GR action in non-neuronal cells influences memory recall, we combined auditory fear conditioning in mice and the knockout of GRs in astrocytes in the prefrontal cortex (PFC), a brain region implicated in memory recall. We found that GRs in astrocytes in the PFC calibrate recall in female but not male mice. Specifically, we found that knocking out GRs in astrocytes in the PFC of female mice (AstroGRKO) after fear conditioning resulted in higher recall of fear to the CS+ tone when compared to controls (AstroGRintact). While we did not find any differences in extinction of fear toward the CS+ between these groups, AstroGRKO female mice showed impaired recall of extinction training. We did not observe any significant results in male mice. These results suggest a sex-specific calibration of memory recall by GRs in astrocytes in the PFC. These data demonstrate the need to examine GR action in cortical astrocytes to elucidate the basic neurobiology underlying memory recall and potential mechanisms that underlie female-specific biases in the incidence of PTSD.

## INTRODUCTION

Recalling important information about salient environmental cues is an integral part of how we navigate our world. Recalling too much, or too little, information about salient environmental cues is a part of the psychopathology of Post-Traumatic Stress Disorder (PTSD). More specifically, the augmented recall of an association between an environmental cue and a traumatic event results in a debilitating fear toward the cue, even in the absence of any threat. In contrast, an impoverished recall of information that a cue, previously associated with trauma, is no longer a threat results in debilitating fear toward the cue after it is no longer dangerous. Therefore, one way to mitigate debilitating fear that characterizes PTSD is to understand the neurobiological mechanisms underlying both, the enhanced recall of fear toward cues that have been previously associated with an aversive outcome and the diminished recall of information that this same cue is no longer threatening.

Among many mechanisms, glucocorticoid action via signaling through glucocorticoid receptors (GRs) is an important neurobiological pathway that underlies the recall of salient information. When trauma-associated cues are encountered the hypothalamic-pituitary-adrenal axis is activated and GR signaling is consequently triggered [1–3]. Existing literature demonstrates that glucocorticoids and GRs do in fact influence learning, memory and the recall of learning [4–32]. As expansive as this research is, the influence of GRs on learning, memory and recall of learning has mostly focused only on GR action in neurons or has ignored cell type specific contributions all together. While glia are approximately as common as neurons in the nervous system [33,34], the role of GRs in glial cells on the recall of salient environmental cues has been neglected. More specifically, while astrocytes represent 19-40% of glial cells [33] and express GRs [35,36], the influence of GRs in astrocytes on memory recall remains largely unappreciated (for one exception, see Discussion).

Our goal in this study was to determine the influence of GRs in astrocytes on memory recall. To do so, we combined the robust and reliable experimental framework of classical fear conditioning in rodents [37–44] with molecular genetic manipulations in the prefrontal cortex (PFC), a brain region critical for the recall of memory [45–51]. We first trained mice to associate tone presentations with mild foot-shocks. After this auditory fear conditioning, we used a CRE-loxP strategy to specifically knock-out GRs in astrocytes in the PFC of these trained mice. We then exposed animals to extinction training: tones in the absence of any foot-shocks. This allowed us to ask how a lack of GRs in astrocytes in the PFC (i) influences the recall of the previous aversive association of the tone presentation with the foot-shock, and (ii) influences the acquisition of extinction of fear responses that would typically occur during extinction training. Finally, we exposed animals to tone presentations one day after extinction training, and this allowed us to measure the influence of GRs in astrocytes in the PFC on the recall of extinction training. Broadly speaking, our results demonstrate that GRs in astrocytes in the PFC are required for the normative recall of memories in female but not male mice: the recall of memory associated with the conditioning event, as well as the recall of extinction training.

## MATERIALS AND METHODS

### Animals

Nr3c1-floxed (GR-floxed) animals were obtained from Jackson Laboratories (Strain #: 012914) and then back-crossed to a C57BL/6J background. Homozygous GR-floxed and wild-type littermate controls were group-housed in the vivarium located in the Neuroscience Building of the Yerkes National Primate Research Center, with controlled temperature, humidity and pressure, kept on a 14:10-h light/dark cycle, and given *ad libitum* access to food and water. All experiments were approved by the Emory Institutional Animal Care and Use Committee and followed NIH standards.

### Auditory Fear Conditioning

Behavioral sessions were conducted in conditioning chambers (Coulbourn Instruments) connected to tone and shock generators controlled by FreezeFrame software (Actimetrics). On day 1, animals were habituated to the conditioning chambers by placing them in context A for 5 minutes. Providing a distinct context for fear conditioning, Context A consisted of the chamber with metal rod floors, quatricide as the cleaning agent, and chamber as well as room lights turned off. Infrared lights in the chambers were left on to allow for recording. On day 2, animals were conditioned in context A, as follows: after an acclimation period that lasted 180s, animals presented with five 30-second tones (6kHz, 70dB), termed the conditioned stimulus (CS+). Each tone presentation co-terminated with a 1s, 0.5 mA foot-shock [unconditioned stimulus (US)]. Tone-shock pairings were presented with an inter-trial interval (ITI) that averaged 2 minutes. After conditioning, animals were returned to the vivarium for 3-5 days before intra-cranial surgeries were performed on them as noted below.

### Intra-cranial surgeries

To knock-out GRs in astrocytes of the prefrontal cortex (PFC), GR-floxed mice and littermate controls were injected with an AAV5-GFAP-GFP-Cre virus. Additionally, two control females used were of the GR-floxed genotype and injected with GFAP-GFP expressing virus, leaving GRs intact. These injections resulted in two groups of mice; those in which GRs were left intact in astrocytes in the PFC (AstroGRintact) and those in which GRs were knocked-out in astrocytes in the PFC (AstroGRKO). Viruses were purchased from Addgene. Animals were given oral Metacam (meloxicam) as an analgesic and then anesthetized using a mixture of ketamine and dexdomitor, injected interperitoneally. Animals were mounted into stereotaxic frames (Stoelting) and stereotaxic injections were performed bilaterally into the PFC at the following coordinates relative to Bregma: AP +1.69 mm, ML ±0.16 mm, DV −2.85 mm. Viruses were injected using a Nanoject III (Drummond Scientific) with a pulled glass micropipette. The final volume of AAV-containing solution was 80 nl, injected at a rate of 1 nl/sec. Following injections, the glass pipettes were left in place for 5 minutes before slowly being removed over 1 minute. The scalp was closed with sutures and the animal was administered Antisedan, to counteract the anesthesia. Animals were moved back to the vivarium to recover. Further behavioral testing was conducted two weeks later to allow for optimal expression of the CRE recombinase and/or fluorescent protein.

### Testing recall of fear memory and memory of extinction training

After viral expression (2 weeks post-surgery), animals’ recall of fear memory and of extinction training was measured as follows. Briefly, after 180 secs of acclimation to the same chambers as conditioning, but with distinct contextual indicators (context B), animals were exposed to 30 CS+ tone presentations with 30 secs ITI between CS+ presentations. Specifically, the chambers were fitted with opaque plexiglass floors, both the house lights and room lights were on, and 70% ethanol was used to clean the chambers. Recall of fear memory was measured by averaging freezing during the first 2 CS+ tone presentations. Within-session extinction was measured as the freezing responses to the CS+ presentations in bins of 5 across the entire 30 presentations of the CS+ tones. One day later, recall of extinction memory was measured by presenting 30 CS+ tones in the same manner as extinction training, i.e. 30 CS+ tone presentations in context B.

### Behavioral analyses

All behavior was video recorded and freezing behavior was measured using FreezeFrame-4 software (Actimetrics). The total amount of time spent freezing (in seconds) to the tones was analyzed using FreezeFrame software by an experimenter blind to the treatment conditions.

### Histology

To confirm the expression and placement of intra-cranial virus injections, animals were anesthetized and transcardially perfused using a 4% paraformaldehyde/phosphate-buffered saline solution (PFA/PBS). Brains were stored overnight in 4% PFA/PBS and then transferred to a 30% sucrose/PBS solution until saturated (3-4 days). Brains were then sectioned at 35μm on a freezing microtome (Leica), stained with Hoechst nuclear stain (1:1000), and mounted with SlowFade Gold Antifade mounting solution (Life Technologies). Location of the GFP fluorescent tag was observed with a Nikon Eclipse E800 fluorescent microscope and used to determine virus placement and expression.

### Fluorescence Activated Cell Sorting (FACS) and RT-qPCR to determine GR knock-out in astrocytes

#### Animal brain dissociation

AstroGRintact and AstroGRKO mice were sacrificed after isoflurane anesthesia and whole brains were isolated. Brains was blocked to isolate an approximately 300 um brain section that contained the prefrontal cortex (PFC). The PFC was punched using a 1mm stainless steel tissue punch. The Papain Dissociation Kit (Cat# LK003150 Worthington) was used to dissociate the tissue. Briefly, tissue was minced 100x on glass slide and then added to 1 mL of Worthington papain solution. The tissue was then placed at 37°C in a shaking incubator for 30 minutes and triturated every 5 minutes with a fire polished 5ml glass Pasteur pipet. Un-dissociated tissue was allowed to settle, and supernatant was placed in a new tube and centrifuged 4°C for 5 minutes at 300g. The cell pellet was suspended in 1 ml Worthington albumin-ovomucoid inhibitor solution 4°C for 5 minutes. The resulting solution was centrifuged for 5 minutes at 300g at 4°C and the pellet suspended in .5 ml PBS 1% FBS FACS solution.

#### FACS Sorting

Dissociated brain cells were filtered into FACS tubes with filter tops (Cat# 352235 BD) to eliminate clumps and then FACS sorted using a FACS ARIA II FACS sorter. Cells were gated using forward scatter (FSC-A) and side scatter (SSC-A). Clumps and doublets were removed by gating singlets in two linear scale dot-plots of SSC-W versus SSC-H and FSC-W and FSC-H. GFP positive cells were sorted based on fluorescence intensity and size of cells and gated. Approximately 20,000 cells were collected at 4°C in 1.5 ml micro-centrifuge tubes containing PBS 1% FBS FACS solution. Cells were pelleted for RNA Extraction.

#### RNA extraction and qRT-PCR

RNA for the FACS sorted cells was extracted using the Pico Pure RNA Isolation kit (Cat#kit0204 Arcturus). Purification was performed according to the manufacturer’s protocol using RNA extraction from cell pellet protocol with on-column Dnase treatment and eluted with 12 ul elution buffer. cDNA was synthesized from 40 ng of isolated RNA using the RT2 First strand Kit (Cat#330404 Qiagen). Pre-Amplification of cDNA was done using Single cell to CT kit (Cat#445 8237 Thermo Fisher Scientific) and Taqman gene expression assay primer sets [gapdh (Cat# Mm99999915_g1), rbfox3/neun (Cat# Mm01248771_m1), gfap (Cat# Mm01253033_m1), nr3c1 (Cat# Mm00433832_m1)]. Standard qRT-PCR was done using Sybr Taqman 2x universal mix (Cat# 430 4437 ABI) with same primer sets on an Applied Biosystem 7500 Fast Real Time PCR system. The 2ddCT method was used to quantify RNA expression.

### Statistics

Statistics were performed with GraphPad Prism. Repeated-measures two-way ANOVA (GR status and CS) was used for behavioral analyses across sessions that consisted of the 30 CS+ presentations and unpaired t-tests were performed where comparisons were explicitly made between two groups at specific timepoints (e.g. recall of fear memory and recall of extinction memory). Post hoc tests were performed for significant interactions. Fishers LSD was used for post-hoc comparisons and p < 0.05 was considered significant.

## RESULTS

### GRs in astrocytes in the PFC are knocked-out using a Cre-LoxP strategy

To knock-out GRs in astrocytes in the PFC, we infused either GFAP promoter driven GFP expressing AAVs or GFAP promoter driven Cre+GFP expressing transcripts into the PFC of GR-floxed mice (Fig. 1A and 1B). This strategy resulted either in no perturbation of the GR receptor (AstroGRintact) or Cre recombinase-mediated knock-out of the GR gene in astrocytes in the PFC (AstroGRKO), as measured by RT-qPCR after extracting RNA from GFP-expressing astrocytes via FACS sorting. More specifically, while GR transcripts were detected in astrocytes of the AstroGRintact group, no GR transcripts were detected in astrocytes of the AstroGRKO mice at the end of the 40 cycles of qPCR (Fig. 1C). Equal amounts of GAPDH transcripts were detected in the AstroGRintact and AstroGRKO groups (Fig. 1C) (n = 4/group, ANOVA: F(3,12)=63.44, p < 0.0001. Post-hoc comparisons: AstroGRintact GAPDH Ct vs. AstroGRKO GAPDH Ct, p > 0.05, AstroGRintact GR Ct vs. AstroGRKO GR Ct p < 0.0001).

**Figure 1:**
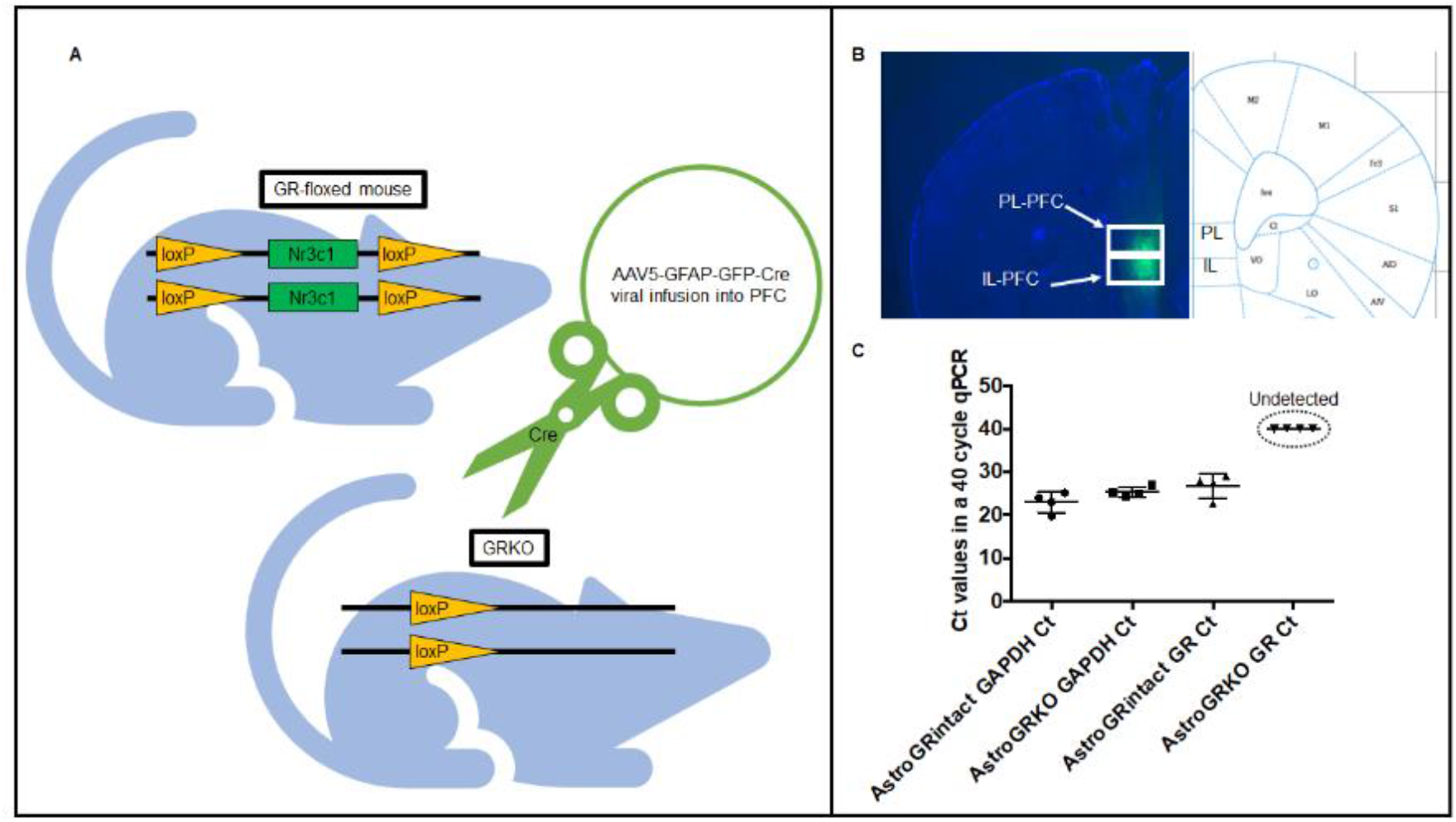
Viral infusions into the PFC knock out GRs in astrocytes. **(A)** An AAV5-GFAP-GFP-Cre virus was injected into the PFC of GR-floxed mice to induce Cre recombinase-mediated knock-out of the GR gene in astrocytes in the PFC. **(B)** Representative image of PFC injections which span both the prelimbic (PL) and infralimbic (IL) sub-divisions of the PFC (Allen Mouse Brain Atlas). **(C)** Astrocytes were isolated using FACS and qPCR was performed to determine GR gene expression in astrocytes. Transcripts of GAPDH were detectable in both AstroGRKO animals (Ct = 25.40 +/- 0.5903) and AstroGRIntact animals (Ct = 23.0 +/- 1.184). Ct values for GAPDH did not differ significantly between the groups (p > 0.05). GR transcripts were detectable only in AstroGRintact animals (Ct = 26.77 +/- 1.386). GR transcripts were undetectable in AstroGRKO animals, not reaching fluorescent threshold in 40 cycles (n = 4/group, ANOVA: F(3,12)=63.44, p < 0.0001. Post-hoc comparisons: AstroGRIntact GAPDH Ct vs. AstroGRKO GAPDH Ct, p > 0.05, AstroGRIntact GR Ct vs. AstroGRKO GR Ct p < 0.0001).

### GRs in astrocytes in the PFC calibrate the recall of fear memory in female mice

Prior to GR knock out, all animals underwent auditory fear conditioning in Context A (Fig. 2A). During auditory fear conditioning (CS+ tones paired with foot-shocks), female and male AstroGRKO mice (female: n = 8, male: n = 9) show the same acquisition of freezing to the CS+ (conditioned stimulus) as AstroGRintact mice (female: n = 9, male: n = 6) (Fig. 2B and 2D). The freezing of both female and male mice did not vary significantly as a function of ‘GRStatus’, which was expected as GRs have not yet been knocked out (Repeated measures two-way ANOVA: Female: F(1, 15) = 2.017, p = 0.18, Male: F(1, 13) = 0.01, p = 0.93).

**Figure 2:**
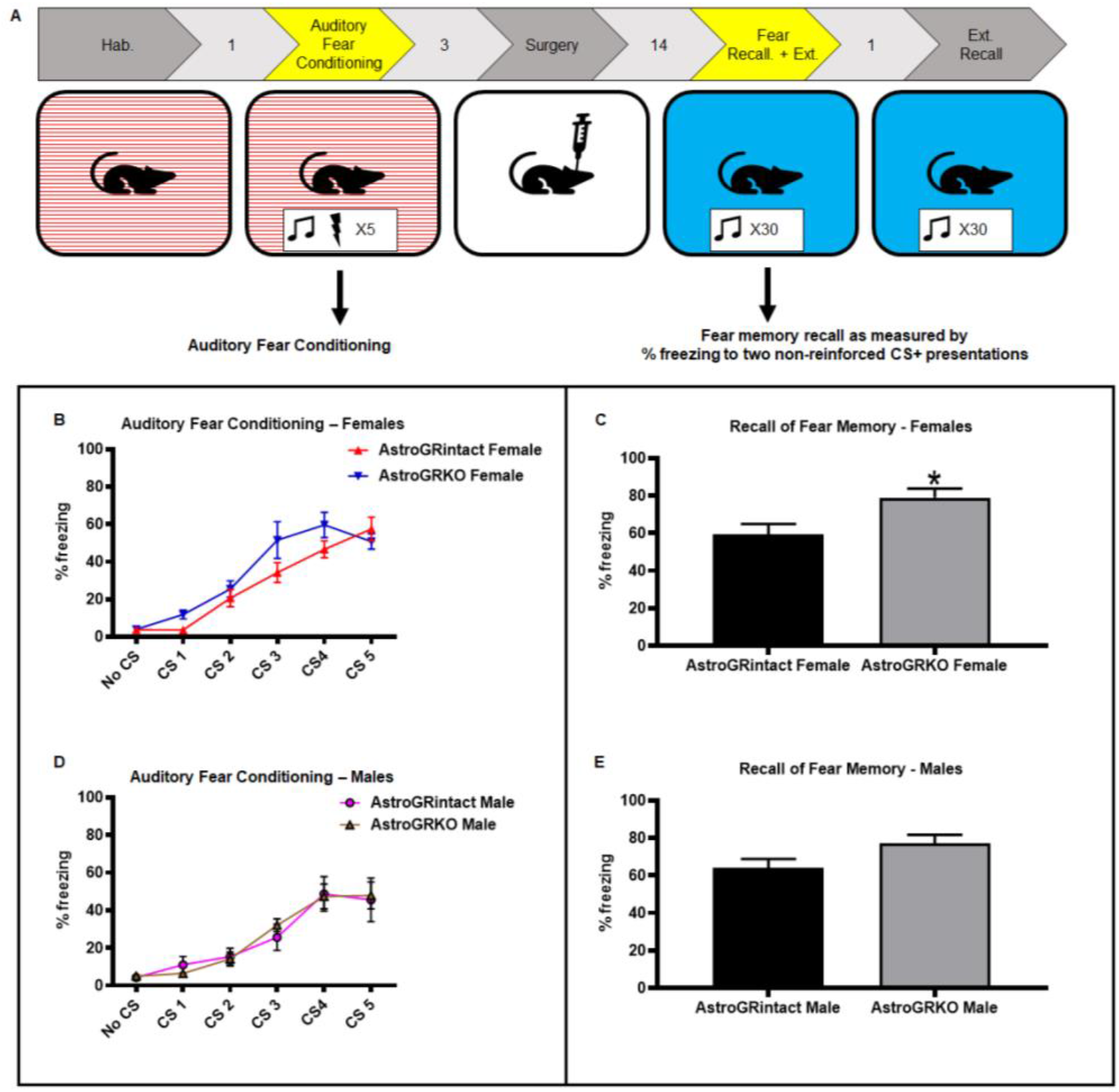
Loss of GRs in astrocytes in the PFC disrupts normative recall of fear memory in female, but not male mice. **(A)** Experimental design showing data in this panel depicting recall of fear memory when GRs are knocked out in astrocytes in the PFC as noted in Fig. 1 after animals have been exposed to auditory fear conditioning. **(B)** Prior to GRs being knocked-out in the PFC of AstroGRKO mice (which occurs after fear conditioning), female mice show no differences in the acquisition of fear during auditory fear conditioning when compared to the AstroGRintact females. **(C)** One day after fear conditioning, AstroGRKO female mice froze significantly more to the CS+ compared to AstroGRintact female mice. (t = 2.596, df = 15, p = 0.02). **(D)** Like seen in the female mice, male mice show no differences in the acquisition of fear during auditory fear conditioning. **(E)** In contrast, one day after fear conditioning, AstroGRKO male mice showed no significant differences in freezing to the CS+ after fear conditioning compared to AstroGRintact male mice (t = 1.838, df = 13, p = 0.089). Data show percent time freezing to the CS+ and are represented as Mean±SEM. Recall of fear memory is defined as the percent freezing during first 2 presentations of CS+ in the extinction training session.

Following auditory fear conditioning, the mice were left to consolidate memory of the conditioning over 3-5 days before receiving intra-cranial injections of AAV-GFAP-GFP or AAV-GFAP-CRE+GFP into the PFC. To ensure adequate expression of GFP or CRE recombinase, animals were left for two weeks and then presented with two, non-reinforced CS+ presentations in Context B (Fig. 2A). Freezing to these two CS+ presentations is reported as a measure of the recall of fear memory. AstroGRKO female mice froze significantly more than AstroGRintact female mice, (Fig. 2C) (Unpaired t-test: t = 2.60, df = 15, p = 0.02). AstroGRKO male mice showed no significant differences in freezing in response to the first two CS+ presentations compared to AstroGRintact male mice (Fig. 2E) (Unpaired t-test: t = 1.84, df = 13, p = 0.09).

### GRs in astrocytes in the PFC do not influence extinction of fear

We found a significant GRstatus x CS interaction when comparing the freezing of AstroGRKO female mice to freezing of AstroGRintact female mice to 30 non-reinforced CS+ presentations in Context B during extinction training (Fig. 3A). As noted above, AstroGRKO female mice froze significantly more to the initial few presentations of the CS+ than AstroGRintact female mice (Fig. 3B) (Repeated Measures Two-Way ANOVA: F (5, 75) = 2.79, p = 0.02. Post-hoc – CS 1-5 p =0.02). However, notably, AstroGRKO female mice showed similar extinction of fear to the CS+ as AstroGRintact mice by the end of the training session. AstroGRKO male mice showed no significant differences in freezing from AstroGRintact male mice at any point in the extinction training that consisted of 30 CS+ presentations in context B with no accompanying US presentations, and both these groups showed similar extinction of fear to the CS+ by the end of the training session (Fig. 3C) (Repeated measures two-way ANOVA: p > 0.05).

**Figure 3:**
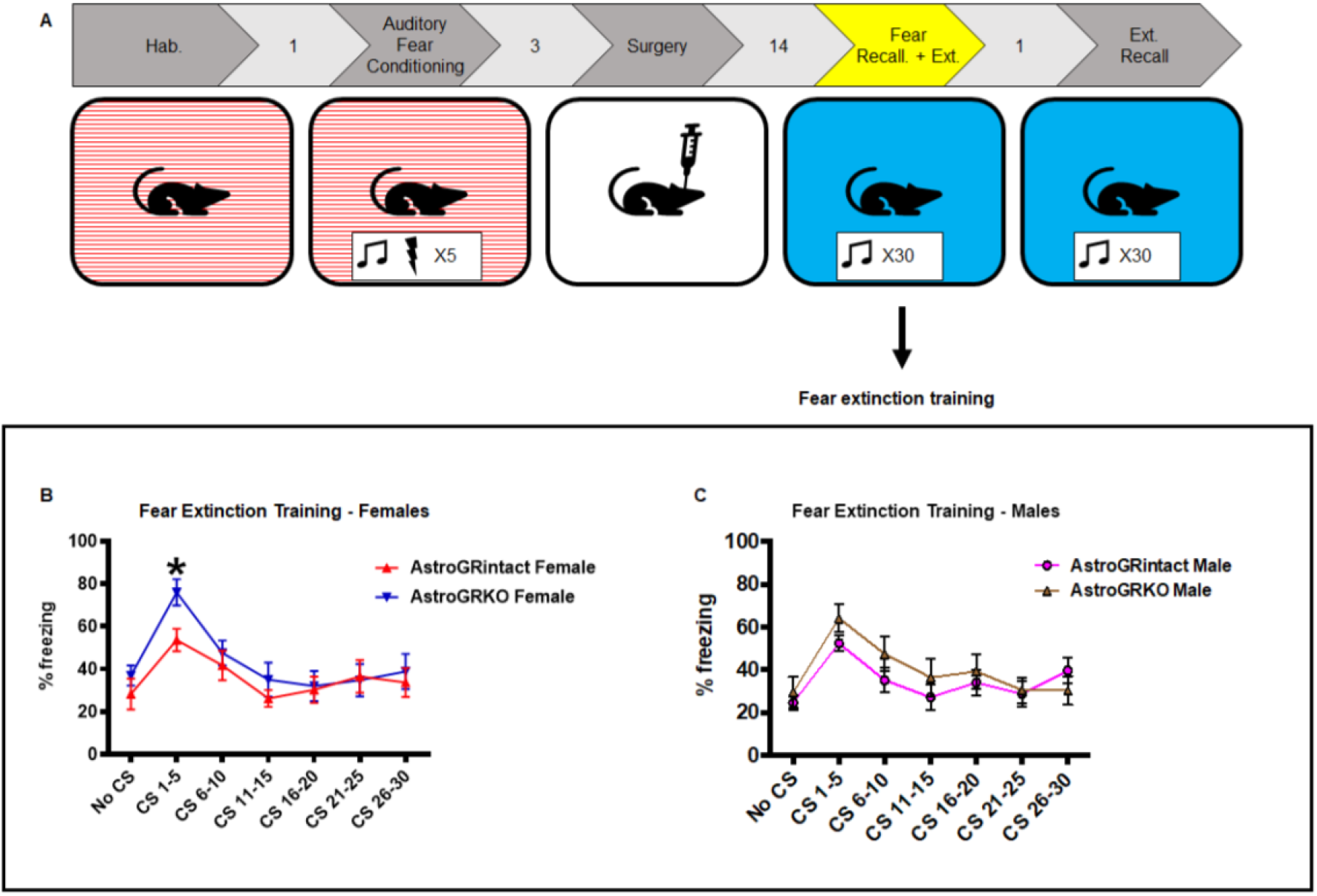
Loss of GRs in astrocytes in the PFC does not impact extinction during extinction training. **(A)** Experimental design showing data in this panel depicting acquisition of extinction during extinction training when GRs are knocked out in astrocytes in the PFC as noted in Fig. 1 after animals have been exposed to auditory fear conditioning. **(B,C)** Following GRs being knocked-out in astrocytes in the PFC after auditory fear conditioning and before exposure to extinction training, AstroGRKO male and female mice show no differences in freezing to the CS+ during repeated non-reinforced presentations of the CS+ during extinction training when compared to the AstroGRintact groups. Data show percent time freezing to the CS+ and are represented as Mean±SEM. (Female - Repeated Measures Two-Way ANOVA: GR status x CS interaction F (5, 75) = 2.79, p = 0.02. Post-hoc – CS 1-5 p =0.02).

### GRs in PFC astrocytes calibrate the recall of extinction memory in female mice

One day after extinction training, the mice were returned to context B and exposed to 30 CS+ presentations without US pairings (Fig. 4A). Similar to the previous test, the first two CS+ presentations in this session were considered as a measure of the recall of extinction training. AstroGRKO female mice showed significantly higher freezing than AstroGRintact female mice to the first two CS+ presentations (Fig. 4B) (Unpaired t-test: t = 2.283, df =15, p = 0.03). Across the entire testing session, we found a significant main effect of “GR status” on freezing to the CS+ presentations between AstroGRKO female mice and AstroGRintact female mice (Fig. 4C) (Repeated measures two-way ANOVA: F (1, 15) = 4.67, p = 0.047). Specifically, AstroGRKO female mice froze more to the CS+ than AstroGRintact female mice during CS+ presentations 1-5 (p = 0.03), 11-15 (p = 0.04) and 16-20 (p = 0.02). AstroGRKO female mice also froze significantly more than AstroGRintact female mice averaged over the full 30 CS+ presentations (Fig. 4D) (Unpaired t-test: t = 2.161, df = 15, p = 0.047). AstroGRKO male mice showed no significant differences in freezing to the first two CS+ presentations from AstroGRintact male mice (Fig. 4E) (Unpaired t-test: t = 1.04, df = 13, p = 0.32). The freezing behavior of AstroGRKO male mice did not vary significantly as a function of ‘GRStatus’ through the entire testing session (Fig. 4F) (Repeated-measures two-way ANOVA: F (1, 13) = 0.16, p = 0.69). Furthermore, AstroGRKO and AstroGRintact male mice showed no significant differences in freezing averaged over the 30 CS+ presentations (Fig. 4G) (Unpaired t-test: t = 0.40, df = 13, p = 0.69).

**Figure 4:**
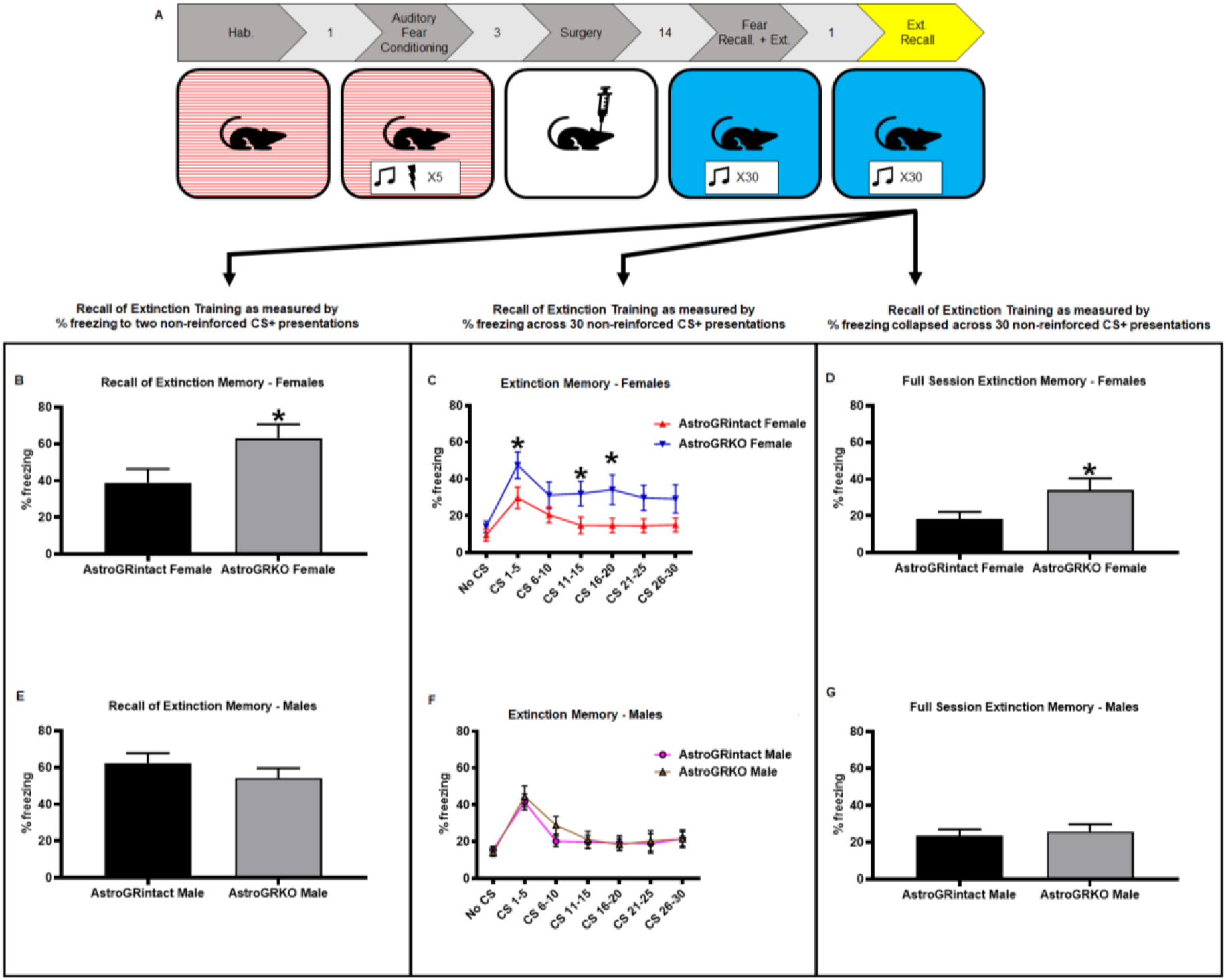
Loss of GRs in astrocytes in the PFC disrupts normative recall of extinction training in female, but not male mice. **(A)** Experimental design showing data in this panel depicting recall of extinction training when GRs are knocked out in astrocytes in the PFC as noted in Fig. 1 before extinction training but after animals have been exposed to auditory fear conditioning. **(B-D)** AstroGRKO female mice showed significant differences in freezing to the CS+, one day after extinction training compared to AstroGRintact female mice. AstroGRKO female mice froze significantly more to the to the first two non-reinforced presentations of the CS+ one day after extinction training compared to AstroGRintact female mice. (t = 2.283, df = 15, p = 0.03) **(B)**. Across the 30 non-reinforced CS+ presentations, one day after extinction training, we found significant main effects for GRstatus (F(1,15) = 4.670, p = 0.04) and CS (F(5,75) = 7.981, p < 0.0001). Post-hoc comparisons indicated that AstroGRKO female mice froze more to the CS+ than AstroGRintact controls in the CS1-5 (p = 0.03), CS 11-15 (p = 0.04) and CS 16-20 (p = 0.02) bins **(C)**. Collapsed across the 30 non-reinforced CS+ presentations, one day after extinction training, AstroGRKO female mice froze significantly more to the CS+ over the full session of extinction memory than AstroGRintact female controls (t = 2.161, df = 15, p = 0.047) **(D)**. **(E-G)** AstroGRKO male mice showed no significant differences in freezing to the CS+, one day after extinction training compared to AstroGRintact male. No significant differences were observed in response to the first two presentations of the CS+ **(E)**, across 30 non-reinforced presentations of the CS+ **(F)**, and collapsed across all 30 non-reinforced presentations of the CS+ **(G)**. Data show percent time freezing to the CS+ and are represented as Mean±SEM. Recall of extinction memory is defined as the percent freezing during first 2 presentations of CS+ in extinction memory testing.

## DISCUSSION

In this study, we combined the robust and reliable experimental framework provided by auditory fear conditioning with molecular genetic-based knock-out of GRs in astrocytes of the mouse PFC to ask whether and how GRs in astrocytes in the PFC impact learning and memory. To do so, we knocked-out GRs in astrocytes in the PFC of female and male mice after CS+ tone+shock fear conditioning. We found that knocking-out GRs in astrocytes in the PFC of female mice after fear conditioning resulted in high levels of fear toward subsequently encountered CS+ tones, and also high levels of fear toward CS+ tones encountered after subsequent extinction training, during which CS+ tones had been presented in the absence of any foot-shock. We did not observe these effects in male mice. These results provide strong evidence that GRs on astrocytes in the PFC play an important role in normative recall of salient environmental events in females, but not males.

Heightened expression of fear toward stimuli previously associated with trauma is a debilitating and highly prevalent dimension of PTSD [52–54]. Such heightened fear could be the result of information that links stimulus and traumatic outcome becoming overly consolidated at the time of this juxtaposition. Alternatively, it could be the result of memory about the association being recalled differently when the stimulus is encountered in the future. In our experimental design, we knocked-out GRs in astrocytes in the PFC several days after the auditory fear conditioning and outside the period during which consolidation of the fear conditioning would occur. This timeline leads us to interpret our finding of AstroGRKO female mice showing increased fear toward the initial presentations of the CS+ tones much after CS+ tone+shock fear conditioning as evidence for GRs in astrocytes in the PFC of female mice being important to calibrate the normative recall of fear. To dissociate the influences of GRs in astrocytes in the PFC on consolidation versus recall, future experiments would need to use as yet unavailable conditional approaches to knock-down these GRs only at the time of consolidation and for GR expression to recover and be at baseline at the time of recall.

Another highly prevalent dimension of PTSD is the expression of fear toward stimuli that may have been previously associated with trauma but are no longer threatening [55–58]. This dimension manifests as heightened fear to the CS+ tones even after extinction training, during which the CS+ tones are no longer paired with a foot-shock. Such debilitating fear could result from an inability to learn that stimuli are no longer threatening, an inability to consolidate this information, or an inability to recall this information. We find that AstroGRKO female mice show within-session extinction that is indistinguishable from controls. These data argue against an influence of GRs in astrocytes in the PFC on learning within an extinction training session that stimuli are no longer threatening. Notably, one day after extinction training, AstroGRKO female mice show increased fear toward CS+ presentations. These findings, while robust, come with the important caveat that they do not allow us to distinguish whether GRs in astrocytes in the PFC impair the consolidation of extinction training or the recall of extinction training. To differentiate these two phenomena, it is again important for future work to use as yet unavailable conditional approaches to knock-out GRs in astrocytes in the PFC only at the time of extinction training, leaving them intact during extinction recall, and vice versa.

It is important to recognize that our data demonstrate the contributions of GRs in astrocytes of the entire PFC to memory recall. Our injections extended dorsoventrally through both the prelimbic-PFC (PL-PFC) and infralimbic-PFC (IL-PFC). These areas are known to have distinguishable and nuanced contributions to learning, memory and fear expression. First, broadly speaking, the PL-PFC is thought to contribute to the expression of fear and the IL-PFC to the suppression of fear [51]. Second, the PL-PFC has been shown to be important for the learning, consolidation and expression of fear associated with the initial conditioning event, while the IL-PFC is more well known for its role in extinction learning and the inhibition of fear after extinction training [39,46,48,49,59,60]. The knockout of GRs in astrocytes of the PL-PFC could be contributing to the differences in the initial expression of fear during recall of the auditory fear conditioning. In contrast, knockout of GRs in astrocytes of the IL-PFC could be mediating the observed effects on consolidation or recall of extinction training. All of the aforementioned work delineating the role of the PL-PFC and IL-PFC focused on the contributions of neurons. It is possible that astrocytes play a distinct role from the neurons in these regions. Future research will have to determine how GRs in astrocytes specifically of the PL-PFC and IL-PFC contribute to memory recall and compare these contributions with those of GRs in neurons in these regions.

While the relationships between GR signaling and learning, and memory and recall are complicated [8,61], there is evidence to suggest that glucocorticoids (GCs) and GR signaling have a positive influence on these phenomena. For example, glucocorticoid administration has been shown to dampen fear recall and impact consolidation of fear extinction, while blocking GC biosynthesis has been shown to impair extinction of a fear memory [12,18]. Micro infusions of GCs into the PL-PFC and IL-PFC improved extinction recall and knocking out GRs on neurons of the PL-PFC impaired extinction recall [15–17]. The only study, to our knowledge, that has specifically addressed the role of GRs on astrocytes in extinction showed that knockdown of GRs on astrocytes throughout the brain reduced freezing in a fearful context [62]. This whole brain knockdown occurred prior to fear conditioning and, as was interpreted by the authors, likely represents an impairment in aversive memory formation rather than improved fear extinction. In keeping with these data, we would interpret our work to be suggestive of GRs in astrocytes in the PFC being facilitatory and not disruptive of memory recall.

Thinking about how GRs in astrocytes might calibrate memory recall, we speculate that relationships between GRs and lactate-mediated signaling deserve experimental scrutiny. Not only is astrocyte-neuron lactate signaling important for learning and memory [63,64], but recent work has also demonstrated that GR knockdown in astrocytes decreases neuronal excitability via reduction of GR-induced lactate release [65,66]. Based on these data, in the future, we propose to test whether the altered memory recall that we report is a consequence of astrocyte GR knockout-induced lactate reduction resulting in reduced neuronal activity in the PFC.

An important dimension of our data are the female-specific effects that we report. We did not find any significant differences in the male mice that we studied using the same experimental design to which the female mice were exposed. We do not have any definitive mechanisms via which GRKO in astrocytes impacts memory recall in female, but not male mice. However, we are heartened by a similar pattern that was recently reported. Specifically, knocking-out GRs, prior to fear training, in CaMKIIa neurons in the PL-PFC of female, but not male rats impaired fear extinction recall and heightened fear expression during training [17]. Females are twice more likely than males to develop PTSD after a traumatic event [67]. The female-specific significant data that we report suggest that GR signaling in astrocytes might be an attractive molecular candidate to probe further as related to the aforementioned sex-bias in the incidence of PTSD. Sex is already known to influence numbers of astrocytes in brain regions linked to stress responsiveness. For example, male rodents have more astrocytes in the medial amygdala and females have more astrocytes in the hippocampus [68–70]. Then there are data to suggest that exposure to stress can alter astrocyte biology. In the PFC, chronic stress decreases expression of astrocyte-markers (GFAP) and induces atrophy of astrocyte processes in males, while increasing GFAP expression and complexity of astrocyte anatomy in females [71,72]. Such data demonstrate that characteristics of astrocytes at baseline, expression of astrocytic markers, and stress-induced changes in astrocytes are different in males and females. Therefore, it stands to reason that manipulating GRs in astrocytes will differentially change fear expression in the sexes. Motivated by such literature, future experiments will need to compare and contrast the relative contributions of GRs in astrocytes versus GRs in neurons of the PFC and continue focusing on sex as a key biological variable that influences astrocyte biology.

In conclusion, our data provide compelling evidence that GRs on astrocytes are necessary for normative memory recall. To our knowledge, ours is the first report linking GR function specifically in astrocytes in the PFC to fear-related memory recall in a sex-specific manner.

## ACKNOWLEDGEMENTS

We thank the Veterinary and Animal Care staff in the Yerkes Neuroscience Vivarium for animal husbandry.

## AUTHOR CONTRIBUTIONS

BGD conceptualized the project, designed the study, interpreted data and directed manuscript writing. WWT contributed intellectually to the project, performed experiments, analyzed data, interpreted results and wrote the manuscript. BRI, ZSS contributed intellectually to the project, performed experiments, analyzed data and interpreted results. KMG contributed intellectually to the project, was involved in pilot experiments and helped edit the manuscript.

